# Effects of linker length on phase separation: lessons from the Rubisco-EPYC1 system of the algal pyrenoid

**DOI:** 10.1101/2023.06.11.544494

**Authors:** Trevor GrandPre, Yaojun Zhang, Andrew G. T. Pyo, Benjamin Weiner, Je-Luen Li, Martin C. Jonikas, Ned S. Wingreen

## Abstract

Biomolecular condensates are membraneless organelles formed via phase separation of macromolecules, typically consisting of bond-forming “stickers” connected by flexible “linkers”. Linkers have diverse roles, such as occupying space and facilitating interactions. To understand how linker length relative to other lengths affects condensation, we focus on the pyrenoid, which enhances photosynthesis in green algae. Specifically, we apply coarse-grained simulations and analytical theory to the pyrenoid proteins of *Chlamydomonas reinhardtii*: the rigid holoenzyme Rubisco and its flexible partner EPYC1. Remarkably, halving EPYC1 linker lengths decreases critical concentrations by ten-fold. We attribute this difference to the molecular “fit” between EPYC1 and Rubisco. Varying Rubisco sticker locations reveals that the native sites yield the poorest fit, thus optimizing phase separation. Surprisingly, shorter linkers mediate a transition to a gas of rods as Rubisco stickers approach the poles. These findings illustrate how intrinsically disordered proteins affect phase separation through the interplay of molecular length scales.

## Introduction

Biomolecular condensates – organelles without membranes – are used by cells to organize and orchestrate a multiplicity of processes, ranging from signaling^1–3^ and gene regulation^4, 5^ to metabolism^6^ and photosynthesis^7^. The physical properties of these condensates and thus their functions are sensitive to the microscopic features of their constituent biomolecules. Importantly, these microscopic parameters are subject both to evolutionary adaptation via mutation of molecular sequences and to active regulation through chemical modifications. While the interactions that lead to biomolecular phase separation are typically complex, the phase-separated algal organelle called the pyrenoid, which is the locus of carbon fixation, presents a tractable model system in which the interactions are well understood^8–11^. In the leading model alga, *Chlamydomonas reinhardtii*, the multivalency of the two dominant components of the pyrenoid, the rigid Rubisco holoenzyme and its disordered protein EPYC1, are sufficient to drive phase separation in vitro, with each Rubisco binding multiple EPYC1s and vice versa^9^. In other Rubisco condensates such as carboxysomes found in alpha and beta cyanobacteria^7, 13, 14^, other multivalent linker proteins mediate phase separation and the associated stickers on Rubisco are at different locations than in *C. reinhardtii*. Here, we focus on the question of how the microscopic “fit” between EPYC1 and Rubisco in the latter influences phase separation, as a model for the sensitivity of macroscopic phases to microscopic parameters.

The interacting domains of phase separating biomolecules (aka “stickers”) are typically separated by intrinsically disordered regions (IDRs, aka “spacers” or “linkers”). While much attention has been focused on which domains or residues constitute stickers, the linkers also play key roles in phase separation. For example, in experiments the lengths of linkers and their charge content can substantially affect phase separation^15–17^, and influence material coefficients such as viscosity^18, 19^ and viscoelasticity^20^. Linkers also play functional roles, e.g. by recruiting clients such as kinases^21^ and other cargo molecules^22^.

The influence of linker properties on phase separation is confirmed by theory^23–27^. In particular, models with explicit linkers suggest that the volume occupied by linkers within condensates can inhibit phase separation^28^, with a linker’s effective volume depending on its solubility^23, 24^. The flexibility of linkers can also affect phase separation, as seen in models of linker-colloid systems^29–33^. The role of linkers can be modeled implicitly via an effective spring-like interaction between adjacent stickers domains^34–36^. For example, an implicit linker model of RNA has shown that phase separation depends on the effective length of linkers due to the loop-entropy cost of forming sticker bonds^37^.

A remaining open question is how the length of linkers compared to other relevant length scales may affect phase separation. To address this question, we focus on the Rubisco-EPYC1 system, in which the separation between stickers on the rigid Rubiscos is comparable to the effective linker length between EPYC1 stickers. First, we develop a coarse-grained molecular-dynamics model for Rubisco-EPYC1 phase separation based on experimental measurements of size, sticker number and location, and dissociation constant. We find that EPYC1 linker length drastically changes the phase diagram, with shorter linkers favoring the condensed phase. Next, guided by an analytical dimer-gel theory, we trace the sensitivity to linker length to the fit between a single EPYC1 and a single Rubisco. We confirm this conclusion by computationally varying the location of Rubisco stickers, finding the actual locations to be nearly optimal for phase separation. Moving the stickers toward the poles can even lead to an alternative phase in which Rubiscos form one-dimensional rods. Our results suggest that the sensitivity of phase separation to the geometry of binding may more generally provide strong selection pressure on the lengths of IDRs.

## Results

Rubisco holoenzymes (hereafter referred to as “Rubiscos”) and the intrinsically disordered protein EPYC1 combine to phase separate as illustrated in Fig. 1a. While the structure of Rubisco is strongly constrained by its function in carbon fixation, it is not clear what constraints may in principle apply to EPYC1. In particular, what governs the flexible linkers connecting EPYC1’s binding motifs? As shown in Fig. 1b, we address this question via simulations and theory with a focus on the interplay of EPYC1 linker length and flexibility, strength of binding interactions, and Rubisco sticker location. These microscopic parameters affect how EPYC1 binds to Rubisco. As shown in Fig. 1c, when EPYC1 is a “good fit”, almost all its stickers can bind to a single Rubisco molecule; by contrast for a “poor fit”, EPYC1 is not able to bind all its stickers to a single Rubisco. We will explore in detail how this difference influences phase separation.

**Figure 1:**
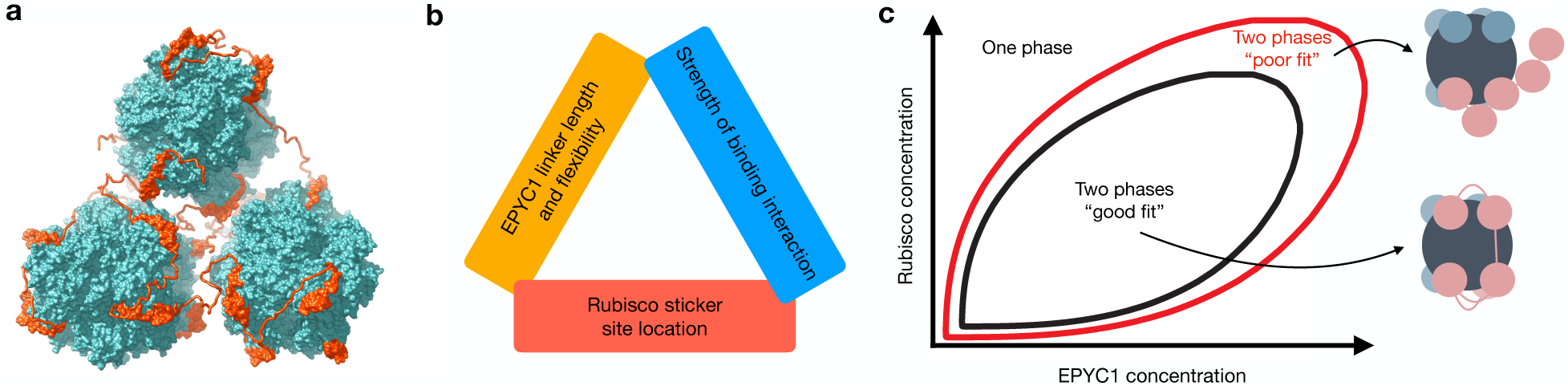
Dependence of Rubisco-EPYC1 phase separation on microscopic aspects. **a** Illustration of three Rubisco holoenzymes held together by several EPYC1s. The Rubisco holoenzymes and EPYC1 sticker regions were obtained from cryo-electron tomography and the EPYC1 linkers were interpolated based on the correct contour length^9, 10^. **b** Three tunable parameters that underpin Rubisco-EPYC1 condensate properties are EPYC1 linker length and flexibility, Rubisco sticker location, and the strength of the binding interaction between EPYC1 and Rubisco stickers. **c** Illustration of how properties of EPYC1 and Rubisco in **b** can control the phase diagram, specifically by giving rise to a “good fit” or a “poor fit” between an EPYC1 and a Rubisco in the heterodimers and other small oligomers that compete with the dense-phase condensate.

We begin our study of the two-component Rubisco-EPYC1 system using coarse-grained molecular-dynamics simulations (Fig. 2). Rubisco is modeled according to its crystal structure^38^ as a sphere with a diameter of about 12 nm, with eight identical stickers symmetrically located near 45 degrees from the top and bottom. EPYC1 is modeled as a polymer of five identical stickers^9^ with an effective linker length between stickers set by a nonlinear spring constant, *k*, which models the entropic free-energy cost of stretching a linker (see Methods for details). All stickers have diameter 𝜎 = 2 nm. Sticker-sticker bonds between EPYC1 and Rubisco are implemented via an associative potential (see Methods). One-to-one bonding is enforced by excluded volume interactions among all other elements.

**Figure 2:**
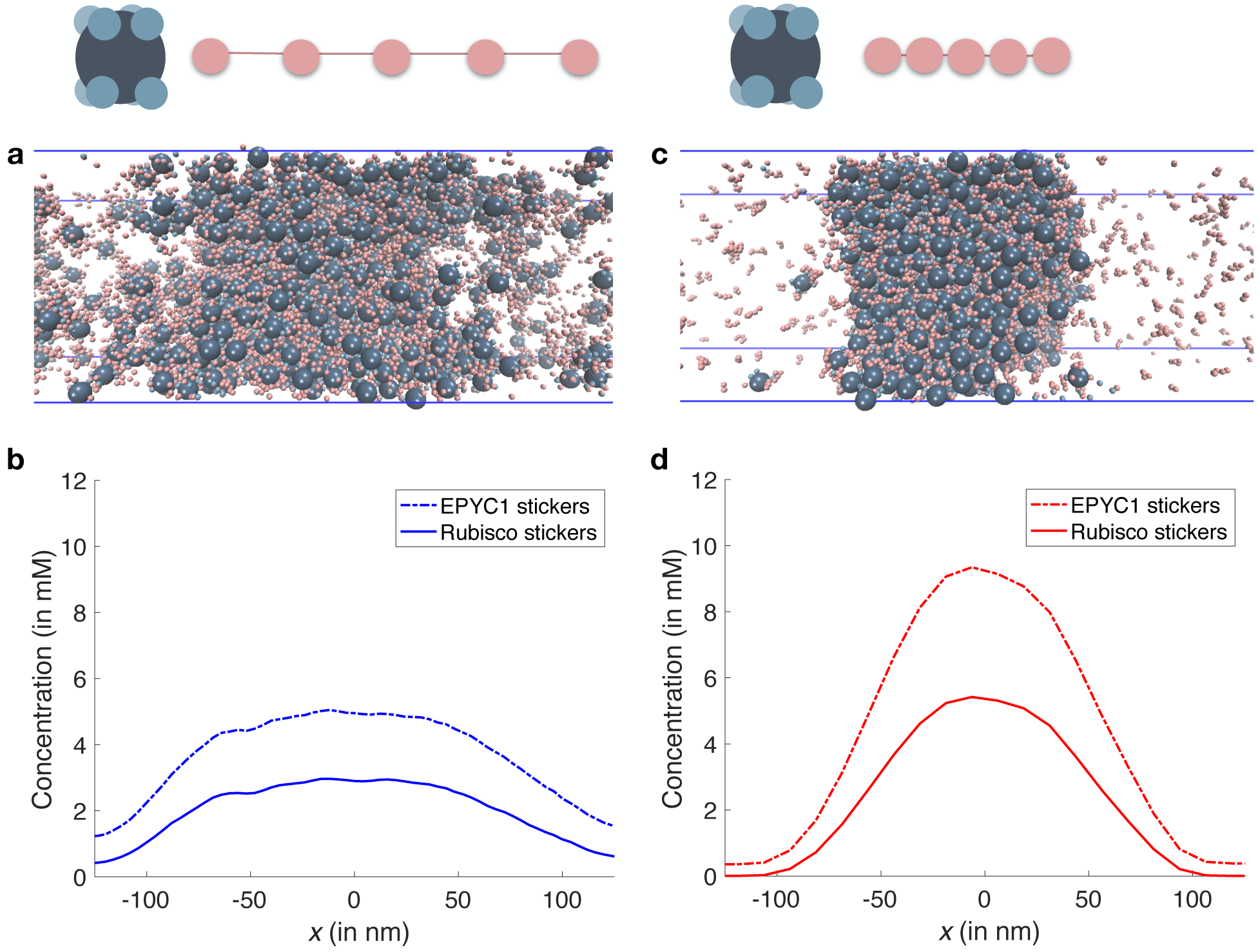
Rubisco-EPYC1 phase separation is sensitive to EPYC1 linker length. As shown in the schematics, the left and right panels show results for EPYC1 linkers, respectively. Data is from simulations of 720 Rubiscos with 8 stickers each and 2,304 EPYC1s with 5 stickers each (i.e. twice as many EPYC1 as Rubisco stickers) in a 500 nm x 126 nm x 126 nm box with periodic boundaries; each sticker has diameter 𝜎 = 2 nm and the sticker binding strength is 𝑈_0_ = 14 𝑘_B_𝑇 (see Methods for details). **a** Snapshot of a phase-separated system with long EPYC1 linkers with a mean equilibrium length of 5 nm (𝑘 = 0.12 𝑘_B_𝑇/𝜎^2^). **b** Sticker concentration profile for EPYC1 (dotted blue) and Rubisco (solid blue) for the system shown in **a** averaged over three simulations. **c** Snapshot of a phase separated system with short EPYC1 linkers with a mean equilibrium length of 2.5 nm (𝑘 = 1.20 𝑘_B_𝑇/𝜎^2^). **d** Sticker concentration profile for EPYC1 (dotted red) and Rubisco (solid red) stickers for the system shown in **c** averaged over three simulations.

Since EPYC1 serves as molecular glue by linking together Rubiscos, it is natural to ask how does EPYC1 linker length affect phase separation given the fixed locations of Rubisco stickers? To address this question, we first employed a long effective EPYC1 linker length of 𝑙 = 5 nm (𝑘 = 0.12 𝑘_B_𝑇/𝜎^2^) which matched the typical sticker-sticker distance from all atom simulations (see Supplementary Note 1). In Fig. 2, we present results of simulations of Rubisco-EPYC1 systems for these long EPYC1 linkers (left column) using Brownian dynamics at a temperature of 𝑇 = 300 K. In Fig. 2a, we show a snapshot of a slab of dense-phase condensate surrounded on both sides by the dilute phase, and in Fig. 2b we show the corresponding EPYC1 and Rubisco sticker density profiles averaged over three independent runs. The total stoichiometry of 2:1 EPYC1 to Rubisco stickers is near the highest imbalance that still produces phase separation for these long linkers (see Supplementary Note 2).

As shown in Fig. 1a, the actual binding geometry of EPYC1 and Rubisco requires EPYC1 stickers to be both compact and oriented in a way that shortens the effective linker length between EPYC1 stickers. We therefore compared the above results for long EPYC1 linkers to those for shorter EPYC1 linkers with an effective EPYC1 linker length of 𝑙 = 2.5 nm (𝑘 = 1.20 𝑘_B_𝑇/𝜎^2^). In Fig. 2c, we show a snapshot for the short linker system, and in Fig. 2d we show the profiles of Rubisco and EPYC1 sticker densities averaged over three independent runs. Comparing the density profiles of the two EPYC1 linker-length systems in Fig. 2b,d, we see that short linkers lead to an almost two-fold increase of both EPYC1 and Rubisco stickers in the condensed phase. In addition, we observe an approximately ten-fold decrease in the critical densities when comparing the long EPYC1 linker system to the short EPYC1 linker system. (Computed by the ratio of the product of the dilute phase concentrations of EPYC1s and Rubiscos for the long and short EPYC1 linker systems.) Moreover, the short linkers lead to phase separation at much higher stoichiometry differences than the long linkers, for example at 10:1 EPYC1 to Rubisco stickers (Supplementary Fig. 2). This implies that the short linkers more strongly favor phase separation than the long linkers under otherwise equivalent conditions.

To better understand the pronounced influence of EPYC1 linker length on phase separation, we next simulated the simple case of a single EPYC1 interacting with a single Rubisco. Specifically, for each of the two different effective linker lengths we obtained the dissociation constant 𝐾_d_ between one EPYC1 and one Rubisco over a range of sticker binding strengths (Fig. 3a). We found that an EPYC1 with long linkers has a lower 𝐾_d_ (stronger binding) to Rubisco since it can form more sticker-sticker bonds with Rubisco than can an EPYC1 with short linkers. As expected, in both cases the dissociation constant 𝐾_d_ decreases exponentially with sticker binding strength. The experimentally estimated dissociation constant between EPYC1 and Rubisco is about 30 nM^11^, which corresponds to 𝑈_0_ = 14 − 15 𝑘_B_𝑇 for the long linkers or around 𝑈_0_ = 18 𝑘_B_𝑇 for the short linkers. For comparison, when 𝑈_0_ = 14 𝑘_B_𝑇 the dissociation constant for EPYC1 with short linkers is ∼2 mM, i.e. almost two orders of magnitude higher than the 𝐾_d_ for long EPYC1 linkers.

**Figure 3:**
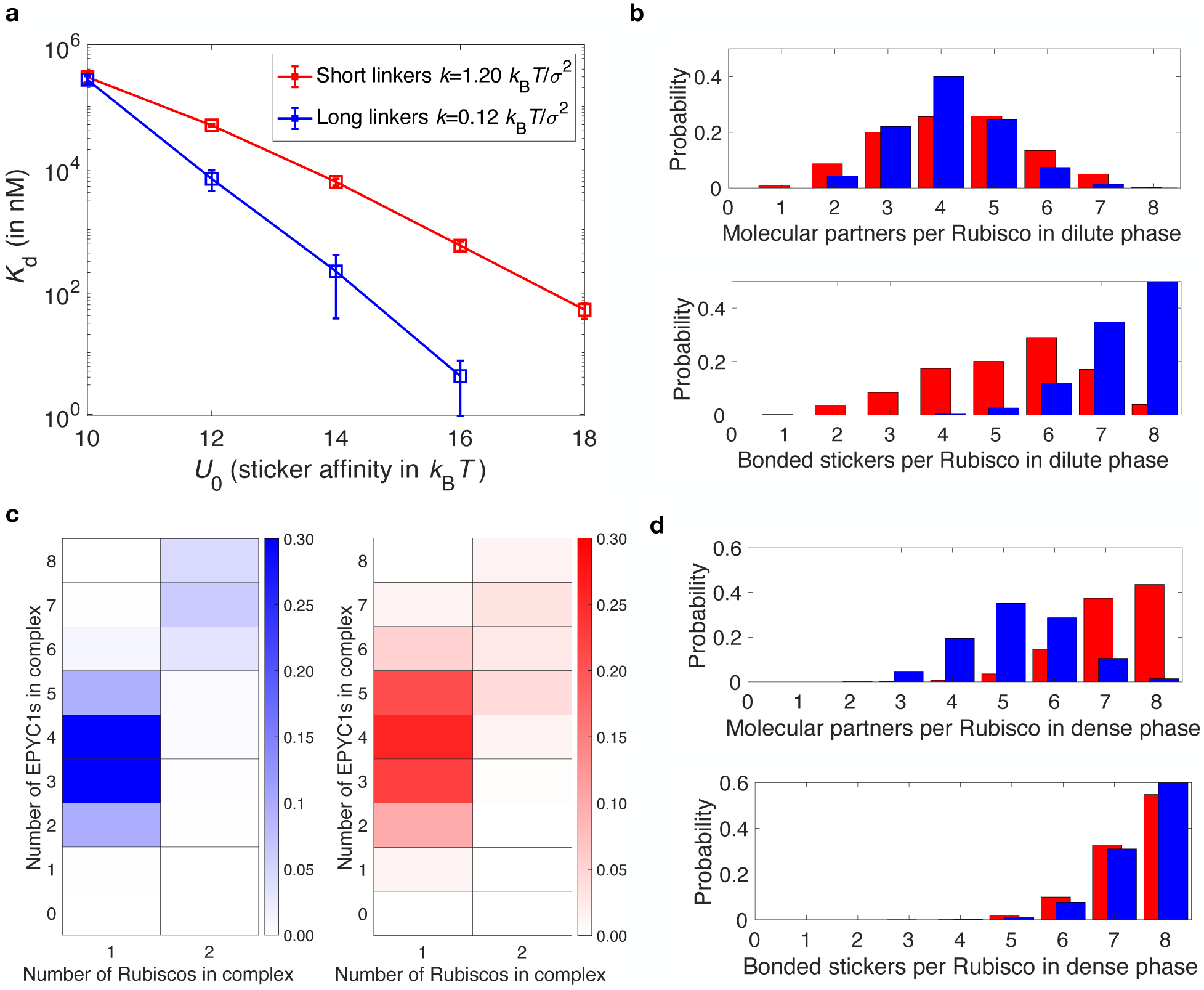
Simulation results for the dilute- and dense-phase properties of Rubisco-EPYC1 systems with short (red) versus long (blue) EPYC1 linkers with the same parameters as in **Fig. 2**.

The large difference in 𝐾_d_ values for different EPYC1 linker lengths begs the question how does this difference in “fit” between one EPYC1 and one Rubisco affect the microscale organization of the dilute and dense phases? Considering first the dilute phase, one potentially informative quantity is how many EPYC1 molecules bind to each Rubisco. As shown in Fig. 3b, we find that these distributions are quite similar for long and short linkers, with a peak around 4 EPYC1 molecules bound to each Rubisco. However, the number of individual Rubisco stickers bound to EPYC1 stickers is quite different: for the long EPYC1 linker system, typically 7-8 Rubisco stickers are bound, whereas for the short linker system, only 4-7 Rubisco stickers are bound. In addition, we asked how Rubisco-EPYC1 association affects the complexes that form in the dilute phase. As shown in Fig. 3c, most complexes had one Rubisco with about 3-4 EPYC1s. For the relatively rare complexes with two Rubiscos, there was typically an additional EPYC1 to bridge them. Next, we consider the dense phase. As shown in Fig. 3d, in sharp contrast to the dilute phase, the typical number of satisfied Rubisco stickers is similarly high ∼7-8 for both long and short EPYC1 linkers. However, the number of EPYC1 molecules bound to each Rubisco is quite different, with 4-6 long-linker EPYC1s versus 7-8 short-linker EPYC1s bound to each Rubisco.

**a** Molecular dissociation constant of Rubisco-EPYC1 dimers as a function of sticker affinity (𝑈_0_) for both short EPYC1 linkers (𝑘 = 1.20 𝑘_B_𝑇/𝜎^2^) and long EPYC1 linkers (𝑘 = 0.12 𝑘_B_𝑇/𝜎^2^). **b** Upper and lower plots show, respectively, the number of EPYC1 molecular partners bound to each Rubisco and the number of bonded stickers on each Rubisco in the dilute phase. **c** Left and right plots show, respectively, the probability of different complexes in the dilute phase. **d** Upper and lower plots show, respectively, the molecular partners bound to each Rubisco and the number of bonded stickers on each Rubisco in the dense phase.

Taken together, these observations suggest an important interplay between EPYC1 linker length and the distance between stickers on a Rubisco. We can gain intuition by considering whether two adjacent EPYC1 stickers are able to bind two adjacent Rubisco stickers. To make this quantitative, we considered a simplified case of a single truncated EPYC1 molecule with only two stickers, one of which is fixed in space, and asked for the probability that the second sticker will be bound to an attractive site a distance 𝑑 away, assuming an implicit linker given by a simple harmonic spring between the two stickers. This probability is given by (see Methods)

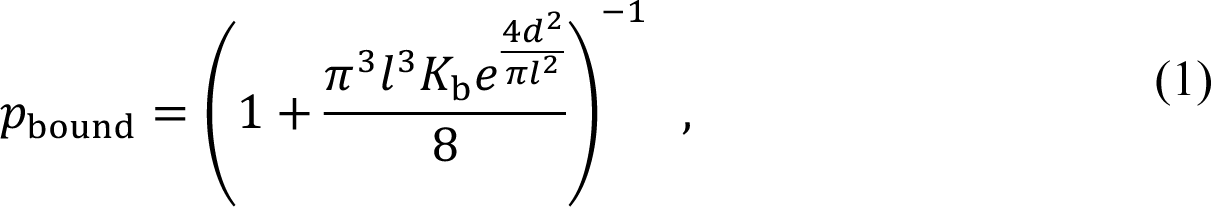

where 𝑙 is the effective linker length, 𝐾_b_ is the dissociation constant of a single EPYC1 sticker from the binding site, and 𝑣 is the effective volume over which the bond can be made. This result demonstrates that there is a large entropic cost to bond consecutive stickers of EPYC1 when the Rubisco sticker separation represented by 𝑑 substantially exceeds the effective linker length 𝑙 between EPYC1 stickers. For a Rubisco sticker spacing of 𝑑 = 6.8 nm, as is the case for Figs. 2 and 3, the exponent 4*d*^2^/(π*l*^2^) in Eq. (1) implies an entropic cost of only ∼1 𝑘_B_𝑇 for a long linker but a much larger cost of ∼10 𝑘_B_𝑇 for a short linker to extend this distance. Hence, for the short linkers it is too entropically costly for neighboring EPYC1 stickers to bind adjacent Rubisco stickers.

This large difference in entropic costs for long and short linkers to bind neighboring stickers on a Rubisco differentially affects the possible configurations of EPYC1s and Rubiscos in both the dilute and dense phases, which in turn influences phase separation. Specifically, the better “fit” between the long EPYC1 linkers and the spacing between Rubisco stickers means that a small number of these EPYC1s can fully satisfy a Rubisco, which allows for energetically favorable small complexes of one Rubisco and 3-5 EPYC1s that compete with the dense phase. By contrast, such favorable complexes are not available for short linkers – either Rubisco stickers are left unsatisfied as shown in Fig. 3b (bottom), or extra EPYC1s would have to be pulled out of solution, with a concomitant translational entropy cost. Instead, the short linkers favor the dense phase within which neighboring EPYC1 stickers can bind to distinct Rubiscos whose stickers can come closer together. Indeed, in the dense phase, short linkers are just as effective as long linkers at satisfying all Rubisco stickers (Fig. 3d, bottom), albeit with each Rubisco sticker bound by a different EPYC1 molecule (Fig. 3d, top). The inverse relationship between molecular fit and phase separation is expanded on further in Supplementary Note 2 where we show that when the Rubisco stickers are allowed to move freely on the main spherical base of each Rubisco there is a greatly improved molecular fit for short-linker EPYC1s, which promotes the formation of dilute phase complexes, and consequently no phase separation is observed at 2:1 EPYC1 to Rubisco sticker stoichiometry. We conclude that the striking differences we observed in our simulations with long versus short EPYC1 linkers can be attributed to the difference in fit between EPYC1 linker length and Rubisco sticker spacing.

It is natural to ask how this difference in fit at the molecular scale influences the overall phase diagram. To this end, we utilize a minimal dimer-gel theory^11, 35^ as shown in Fig. 4. While the model is quite simple, it can capture overall behavior independent of many microscopic details, e.g. the exact types and distribution of dilute-phase oligomers, or dense phase arrangements. The main components of the model are free-energy densities including ideal polymer, excluded volume, and binding contributions (see Methods). In the dilute phase, binding is modeled as formation of small oligomers, which for simplicity we limit to Rubisco-EPYC1 dimers. In the dense phase, binding is modeled as a gas of stickers forming independent sticker-sticker bonds. The only binding-dependent inputs to the model are the dissociation constant 𝐾_d_ between one EPYC1 and one Rubisco and the sticker-sticker dissociation constant 𝐾_b_. For the long and short EPYC1 linkers, the molecular dissociation constants 𝐾_d_ are obtained from Fig. 3a for 𝑈_0_ = 14 𝑘_B_𝑇. The sticker-sticker dissociation constants 𝐾_b_ are chosen to match the dilute-phase concentrations found in the simulations in Fig. 2. Figure 4a shows the overall phase diagrams obtained for these parameters, with a zoomed in version of the dilute-phase boundary shown in Fig. 4b. Consistent with our simulation results, the short EPYC1 linkers strongly favor phase separation, with dilute-phase concentrations as much as an order of magnitude lower than for long linkers. Notably the fitting parameters 𝐾_b_ differ only by 30% compared to the 40-fold difference between the 𝐾_d_ values – thus the striking difference between the phase diagrams can be entirely attributed to the latter. In a nutshell, the better fit between long EPYC1 linkers and adjacent Rubisco stickers allows for stable small complexes in the dilute phase, which disfavors formation of a condensate.

**Figure 4:**
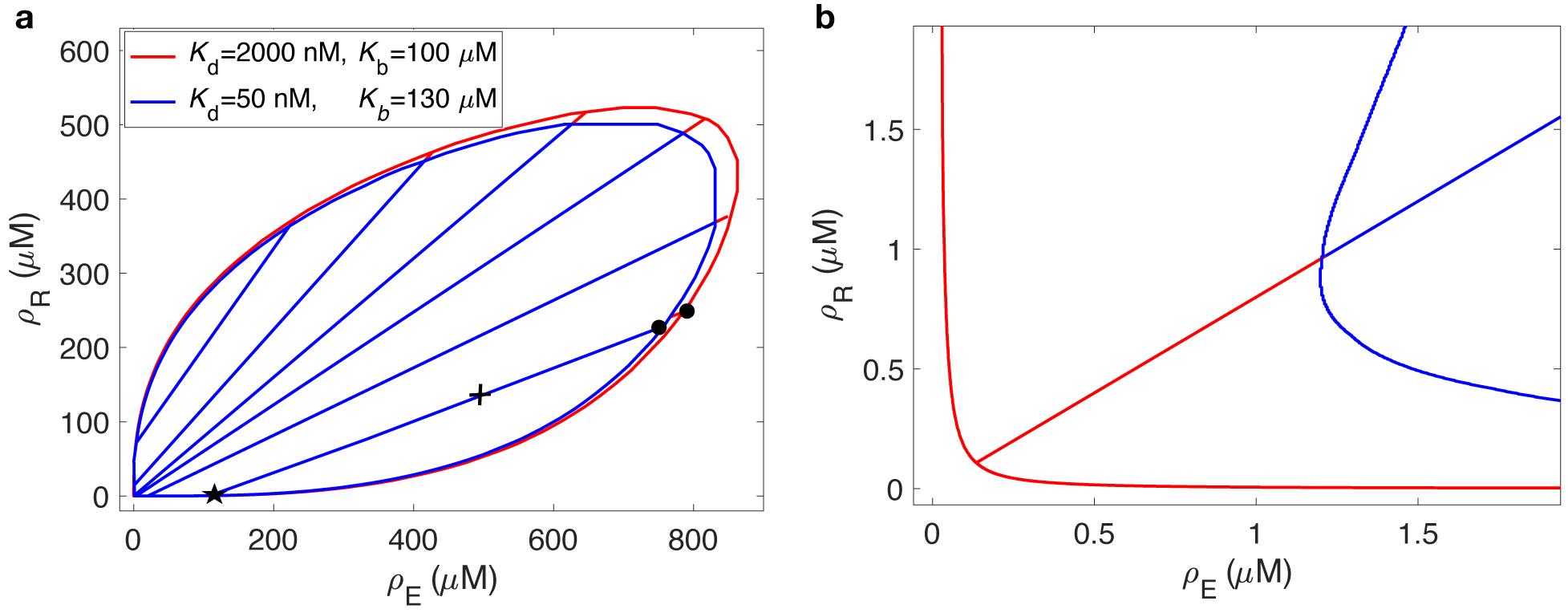
Dimer-gel theory predictions for EPYC1 linker length dependence of Rubisco-EPYC1 phase diagram. **a** Predicted phase diagrams for short (red) and long (blue) EPYC1 linkers using the molecular dissociation constants 𝐾_d_ for 𝑈_0_ = 14 𝑘_B_𝑇. The sticker-sticker dissociation constants 𝐾_b_ are fit so that the dilute-phase concentrations for overall concentrations marked with a “+” agree with the corresponding values from the simulations in Fig. 2. 𝜌_E_ and 𝜌_R_ are the densities of EPYC1 molecules and Rubisco holoenzymes, respectively. Selected tie lines are shown, with the dots indicating dense-phase concentrations for the overall concentrations marked with “+” (the star denotes two overlapping dilute-phase concentrations). **b** Zoom in of the phase diagram in **a** with a representative tie-line (see Methods for details).

Up to this point, we have changed the effective linker length of EPYC1 while keeping the Rubisco stickers at their fixed native locations at 45 degrees from the poles. However, one may ask why has evolution placed the Rubisco stickers of *C. reinhardtii* at particular positions? In Fig. 5, we study the importance of Rubisco sticker location on phase separation. We initially consider the long EPYC1 linkers with mean length 5 nm (𝑘 = 0.12 𝑘_B_𝑇/𝜎^2^). Moving the Rubisco stickers either farther away from the poles to 79 degrees (Fig. 5a) or closer to the poles to 33 degrees (Fig. 5b) both result in weaker phase separation for long linkers. These results are consistent with our above observations that stronger affinity between EPYC1 and Rubisco (lower molecular 𝐾_d_), suppresses phase separation by favoring dimers or other small oligomers of EPYC1 and Rubisco over the condensed phase; as seen in Fig. 5e the 𝐾_d_ values for long linkers are largest near the native Rubisco sticker location at 45 degrees, and decrease as the stickers are moved either farther from or closer to the poles of Rubisco. Thus, the naturally-occurring Rubisco sticker spacing produces the worst molecular fit, but is thereby optimal for phase separation. (Note that changing the sticker-sticker affinity, 𝑈_0_, has little effect on phase separation because, in the language of the dimer-gel theory, it changes both 𝐾_d_ and 𝐾_b_ together, leaving the phase diagram qualitatively unchanged.)

**Figure 5:**
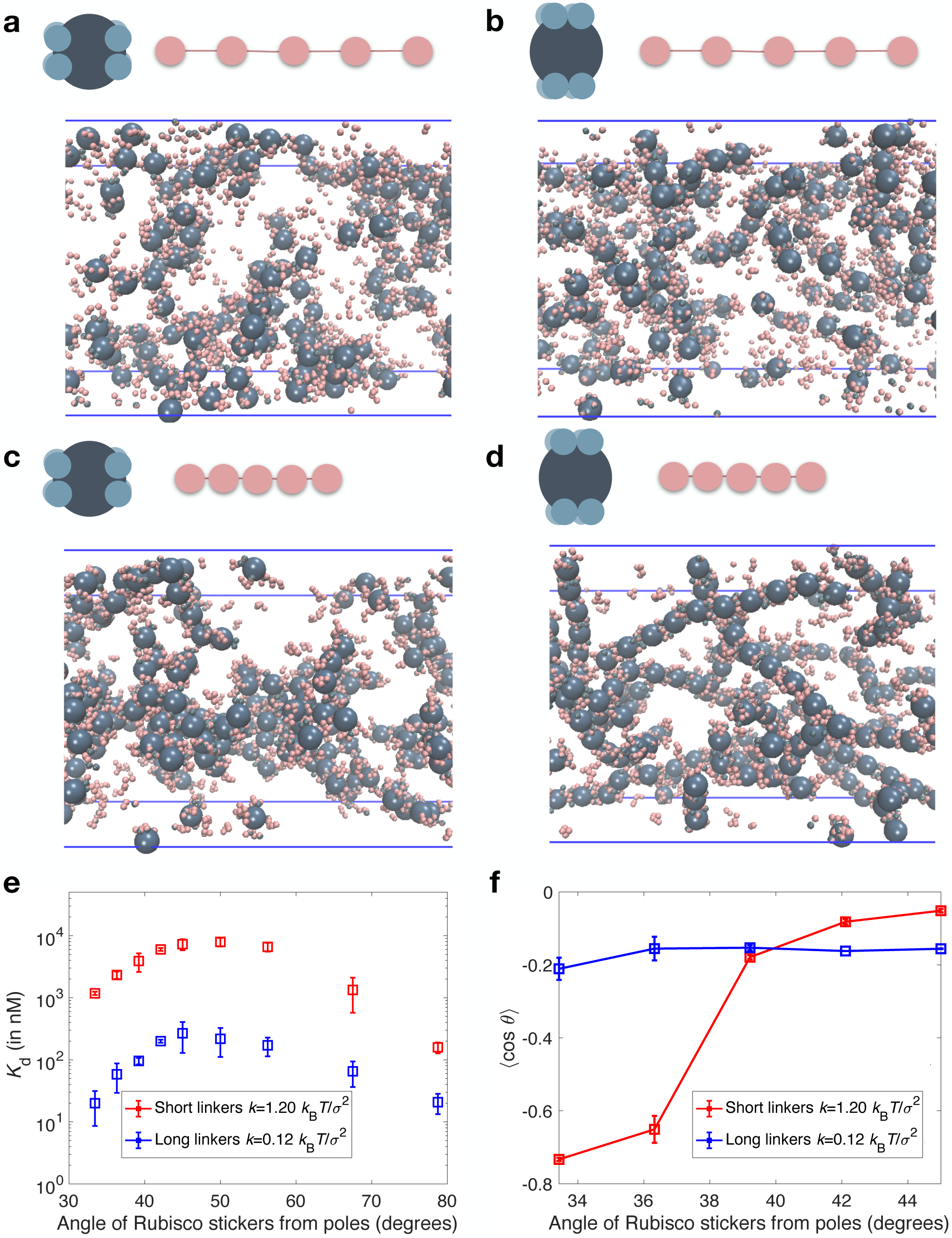
Rubisco sticker location strongly influences phase separation. Rubisco-EPYC1 systems with long (𝑘 = 0.12 𝑘_B_𝑇/𝜎^2^) and short EPYC1 linkers (𝑘 = 1.20 𝑘_B_𝑇/𝜎^2^) were simulated at 2:1 EPYC1 to Rubisco sticker stoichiometry (608 EPYC1s and 380 Rubiscos) for 𝑈_0_ = 14 𝑘_B_𝑇 in a box of size 315 nm x 126 nm x 126 nm with periodic boundaries at a temperature of 𝑇 = 300 K. **a,b** Snapshots of long EPYC1 linkers and Rubiscos whose stickers are located at **a** 79 degrees and **b** 33 degrees from the poles both show weak phase separation. Each system forms a gas of small complexes. **c** Snapshot of short EPYC1 linkers for Rubisco stickers located at 79 degrees from the poles also shows a gas of small complexes. **d** Snapshot of short EPYC1 linkers for Rubisco stickers located at 33 degrees from the poles shows that the system forms a gas rods. **e** The molecular dissociation constant as a function of Rubisco sticker location for short (red) and long (blue) linkers in a box of size 100 nm x 100 nm x 100 nm. **f** Order parameter 〈cos *θ*〉 for rods as a function of angle of Rubisco stickers from poles, where *θ* is the angle formed by neighboring Rubiscos around a central Rubisco (see Methods for details). The blue symbols are for long linkers and the red symbols are for short linkers. Error bars are SD values obtained from three independent simulations.

Motivated by these results, we wondered whether moving the Rubisco stickers closer to the poles, and thus closer together, would improve the fit between Rubisco and a single EPYC1 with short linkers of the mean length 2.5 nm (𝑘 = 1.20 𝑘_B_𝑇/𝜎^2^), thus lowering 𝐾_d_, and disfavoring phase separation. In Fig. 5e, one sees that the molecular fit does improve for short linkers when the Rubisco stickers move closer to the poles. While moving the Rubisco stickers all the way to 33 degrees did indeed abrogate phase separation (Fig. 5d), we were surprised to see that the system organized into a gas of Rubisco “rods”. We quantified this novel organization using as a simple order parameter 〈cos *θ* 〉, i.e. the average cosine of the angle formed by all pairs of neighboring Rubiscos about each central one, averaged over all Rubiscos (see Methods).

Intuitively, this order parameter will have a value of -1 for perfectly aligned rods, and close to zero for dense packings of Rubiscos. For the phase-separated state of short-linker EPYC1s and Rubiscos with stickers at 45 degrees, we find 〈cos *θ* 〉 ≈ 0, consistent with random packing of Rubiscos; in contrast, for the gas of rods that forms for Rubisco stickers at 33 degrees, 〈cos *θ* 〉 ≈ −0.8, implying close to complete rod-like alignment. The reason for rod formation appears to be quite simple: moving the Rubisco stickers to 33 degrees means that two Rubiscos stacked pole to pole can have four extremely close pairs of stickers, which can be easily bridged by EPYC1 linkers. This arrangement is particular favorable for the short EPYC1 linkers, which would otherwise need to be stretched to bind neighboring sites on the same Rubisco (see Supplementary Fig. 3 for the transition to a gas of rods at equal stoichiometry). We have further tested this hypothesis via a toy model in which the Rubisco stickers are allowed to diffuse freely around the Rubisco sphere (Supplementary Fig. 4). We find that Rubisco stickers group together leading to a much higher affinity for short-linker EPYC1 (lower 𝐾_d_), favoring formation of a gas of small oligomers instead of phase separation, but no tendency to form rods.

## Discussion

Biomolecular condensates are typically “network liquids” held together by specific interactions between sticker domains connected by flexible linkers. While linkers play multiple known roles including occupying volume^23, 24^, adding attractive interactions^15–17^, and recruiting clients^21, 22^, the relation between linker length and other intrinsic length scales has been less studied. We explored this topic in the context of a model for the algal pyrenoid, a condensate formed by specific interactions between the intrinsically disordered protein EPYC1 and the rigid holoenzyme Rubisco. Using coarse-grained molecular-dynamics simulations, we found that EPYC1s with shorter linkers led to substantially stronger phase separation. We traced this effect to a worse “fit” between the shortened EPYC1 linkers and the spacing between stickers on a Rubisco: adjacent stickers on short-linker EPYC1s cannot stretch far enough to bind adjacent stickers on Rubisco. By contrast, in the dense phase, adjacent stickers on EPYC1 can bind to distinct neighboring Rubiscos; consequently, short EPYC1 linkers favor the dense phase. Using a minimal dimer-gel theory to predict the full phase diagram, we found that short linkers can lead to phase separation at up to an order of magnitude lower concentrations than longer linkers.

Additionally, by moving Rubisco sticker locations closer to the poles, we found that the native Rubisco sticker spacing is optimal for phase separation, and discovered a novel state in which Rubiscos form a gas of rods. Our study thus highlights how the interplay of linker length with other relevant length scales, in this case Rubisco sticker spacing, can dramatically influence biomolecular phase separation.

We modeled EPYC1 linkers implicitly with an effective length set by a spring constant. This allowed us to isolate the effect of EPYC1 linker length on phase separation. The real Rubisco-EPYC1 system, even in vitro, is certainly more complicated than our simple model: linkers occupy volume and may engage in attractive interactions, while the amino-acid sequences of EPYC1 stickers include charged residues and are not identical, and these stickers bind directionally to Rubisco. However, we do not expect these details to affect our overall conclusion that the phase diagram and the microscale organization of the dense and dilute phases depends sensitively on the fit between EPYC1 linkers and Rubisco stickers. Going beyond the current model would nevertheless be valuable to understand the interplay of the realistic factors noted above. For example, explicit modeling of linkers might reveal if the linker length is tuned to a happy medium – long enough to bridge Rubisco stickers but not so long as to interfere with phase separation by excluded volume. Improvements to make the dimer-gel theory more quantitative are possible as well – our simulations indicate that higher oligomers can dominate over dimers in the dilute phase, and correlations between sticker-sticker bonds in the dense phase may effectively renormalize the density of independent stickers^35^.

The observed sensitive dependence of phase separation on EPYC1 linker length and Rubisco sticker spacing raises the question whether these two lengths may have been jointly optimized by evolution. In the photosynthetic alga *C. reinhardtii*, the pyrenoid is required for efficient carbon capture and so is under strong functional selection. Interestingly, the pyrenoid undergoes complex dynamics during the cell cycle, dissolving into the chloroplast stroma prior to cell division and reforming in the two daughter cells. Hence, evolutionary optimization may not simply favor strong phase separation, but rather may allow for rapid transitions between a phase-separated and a dissolved state, e.g. upon phosphorylation of EPYC1. Moreover, it is essential to the pyrenoid’s function that it remain liquid, as a relatively small number of Rubisco activase proteins must be able to move freely within the pyrenoid matrix to remove inhibitory substrates from Rubisco catalytic sites. Given these multiple constraints, we are hesitant to make concrete statements about the “best” length of the EPYC1 linker, but we will state what functional requirements the real linker would need to satisfy. The illustration in Fig. 1a demonstrates one way in which Rubisco can form the pyrenoid matrix as observed from cryo-electron tomography^9, 10^. One observation is that an EPYC1 linker would need to bind consecutive stickers of the same Rubisco holoenzyme with proper directionality of binding. In Ref. 9 it was found that within the condensate, the distance between the closest EPYC1 stickers of proximal Rubiscos is about 4 nm, which is close to the long linker length. However, the Rubisco-EPYC1 interaction is not completely understood especially considering that residues of EPYC1 are known to be phosphorylated under some conditions^39^. It could be that phosphorylation changes the effective linker length by turning some EPYC1 stickers off, or more generally, by modifying the Rubisco-EPYC1 interaction. Another functional requirement is that Rubisco-EPYC1 condensates are liquid-like with a high degree of mixing as observed in FRAP experiments^10^. Thus, the EPYC1 linker lengths must be long enough to allow some space between Rubiscos to permit the flow of other pyrenoid-localized proteins.

The concept of linker length is not only of interest in the context of the primary condensate-forming linker, EPYC1, but also in the context of other proteins that localize to the condensate and possess multiple Rubisco binding motifs like those of EPYC1. Specific examples in *C. reinhardtii* are Rubisco-binding membrane proteins (RBMPs) which are localized to the membrane tubules that supply CO2 to Rubisco, and StArch Granules Abnormal (SAGA) proteins which are localized to the starch sheath^40^. Both protein families have multiple Rubisco-binding motifs, spaced in some cases by linkers of similar length to those of EPYC1 (∼70 AAs) or about half that length (∼33 AAs). Having sets of longer and shorter linkers may have functional purposes. One possibility is that the longer linkers may permit binding to a single Rubisco, while the shorter linkers may promote bridging between two Rubiscos, or the longer lengths may still favor bridging Rubiscos, but over larger distances. The tendency to bridge Rubiscos would favor localization of RBMP and SAGA proteins with the Rubisco-EPYC1 condensate where many Rubiscos are in close proximity, rather than to Rubiscos dissolved throughout the stroma.

Moreover, bridging configurations would leave open stickers on each Rubisco for EPYC1, allowing the bridged Rubiscos to remain part of the condensate, rather than RBMPs and SAGAs sequestering Rubiscos away from the pyrenoid.

The concept of molecular fit may also apply to linker-Rubisco interactions within alpha^13^ and beta^14^ cyanobacteria. Each Rubisco holoenzyme in alpha-carboxysomes has eight stickers for its flexible partner protein, CsoS2, grouped into four north-south pairs evenly spread around the equator. In the alpha cyanobacteria *Halothiobacillus neapolitanus*, CsoS2 has four stickers^13^ for Rubisco on its N-terminal domain with an approximate linker length (∼50 AAs) of 6.8 nm (see Methods). By contrast, each Rubisco in beta-carboxysomes has only four stickers for its flexible partner CcmM, and these are evenly spread around the equator. In the beta cyanobacteria *Synechococcus elongatus*, CcmM has three stickers^14^ for Rubisco separated by short linkers (∼30 AAs) with an approximate linker length of 5.2 nm (see Methods). Since the diameter of Rubisco is about 12 nm, both the north-south pairs of Rubisco stickers in alpha-carboxysomes and the individual stickers in beta-carboxysomes are separated by about 9.4 nm around the equator (see Methods). This implies that, like the Rubisco-EPYC1 pyrenoid system, the geometry of binding may favor condensates over small oligomers in carboxysomes. Namely, the stickers on carboxysome Rubiscos may be too far apart for a single partner protein, CsoS2 or CcmM, to fully bond to one Rubisco, favoring instead bridging of multiple Rubiscos and so driving phase separation.

The kind of interplay between length scales we explored in the Rubisco-EPYC1 system may be expected as a general feature of multidomain proteins containing intrinsically disordered linkers. Specifically Sorensen et al.^42^ considered two proteins, each with two stickers connected by flexible linkers such that the proteins could form bivalently bound pairs. They found that avidity, i.e. the enhancement of overall binding due to multivalency, increased with decreasing linker length, presumably due to the increase in the effective concentration of the potential binding partners, albeit with a scaling with linker length that suggested additional linker-associated interactions. This interplay of effective linker length with binding constants in multivalent biomolecules is quite ubiquitous^43^ and can be viewed in terms of an effective concentration of stickers set by linker length^44, 45^ relative to sticker affinity. Moreover, the functional importance of the linker length between binding domains is attested to by the conservation of actual^46^ or effective^47^ linker length.

We have shown via simulations and theory that effective linker length relative to another intrinsic length scale can have a drastic effect on phase separation. In the context of Rubisco-EPYC1 condensation, our predictions could be tested by constructing EPYC1 linkers of differing lengths and measuring changes in the phase diagram, as well as probing for microscale features such as Rubisco rods. In live cells, condensates typically form transiently, e.g. in *C. reinhardtii* the pyrenoid dissolves and reforms during cell division. Thus, condensate dynamics are also important for cellular function, and it is known that linker properties can affect dynamical properties such as diffusion constants and viscosity^18, 19^. We hope our results will inspire future investigations of how cells may use linker properties to tune both the steady-state and dynamical properties of biomolecular condensates.

## Methods

### Model details

The package LAMMPS^48^ was used for simulations. In the model, the rigid molecule, Rubisco, has a spherical base with a diameter of 11.6 nm and has 8 stickers of diameter 𝜎 = 2 nm at 45 degrees from the top and bottom and uniformly distributed around the circumference (as shown schematically in Fig. 2). The flexible molecule, EPYC1, is modeled as a linear polymer with five stickers of diameter 𝜎 = 2 nm connected by implicit linkers described by the following nonlinear potential:

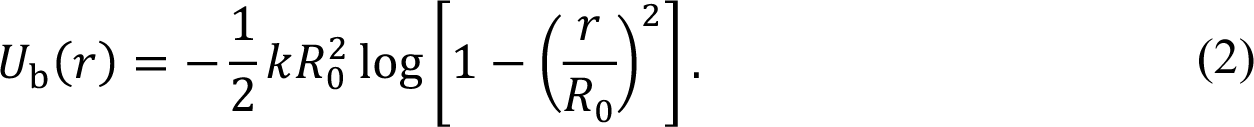

Here, 𝑅_0_ = 19 nm is the length of the linker when fully extended, 𝑟 is the relative distance between two consecutive EPYC1 stickers, and 𝑘 is the spring constant which tunes the effective linker length. The sticker-sticker association between EPYC1 and Rubisco is given by the following potential:

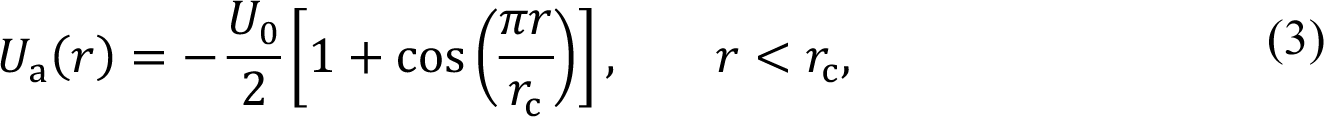

where 𝑈_0_ and the cutoff distance 𝑟_1_ = 𝜎⁄2 = 1 nm define the magnitude and range of the attraction. Additionally, there are excluded volume interactions between all elements of EPYC1 and Rubisco (except stickers of opposite type) given by the potential:

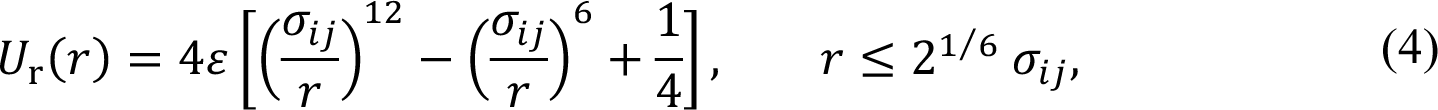

where 𝜀 = 1 𝑘_B_𝑇, 𝜎_*ij*_ is the effective diameter, and 𝑖 and 𝑗 denote the interacting components. For interactions between two EPYC1 stickers or between two Rubisco stickers the effective diameter is 𝜎_EE_ = 𝜎_RR_ = 𝜎 = 2 nm, while for interactions between the Rubisco base and either a Rubisco sticker or an EPYC1 sticker the effective diameter is 𝜎_RE_ = 13.6 nm.

### Dissociation constant **𝑲_𝐝_**

For Fig. 2a of the main text, the molecular dissociation constant was measured in a box of 50 nm x 50 nm x 50 nm with periodic boundaries containing a single EPYC1 and a single Rubisco over a time of 10 ns with a timestep of 1 ps. The EPYC1 and Rubisco are considered to be bound when at least one EPYC1 sticker is within the attractive-interaction cutoff distance 𝑟_1_ of a Rubisco sticker. From two-state kinetics, with one state being the bound state and the other being the unbound state, the dissociation constant is obtained from the following equation:

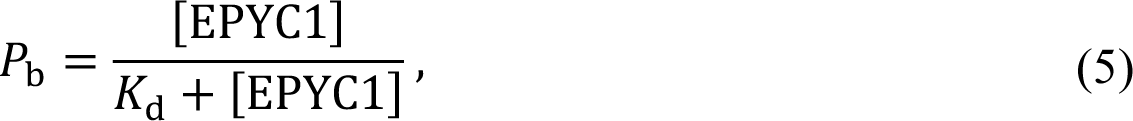

where 𝑃_b_ is the probability, or fraction of time, the two molecules are bound, and the concentration of EPYC1 is given by

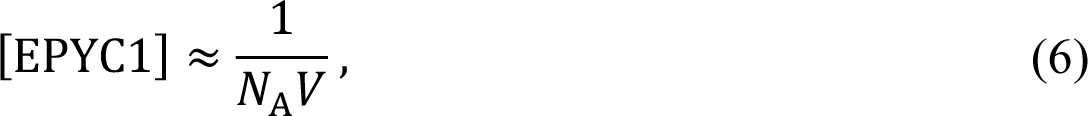

where 𝑁_A_ is Avogadro’s number and 𝑉 is the volume of the simulation box in liters.

### Determination of the dense and dilute phases, satisfied bonds, molecular partners, and complexes

In Fig. 3 of the main text, properties of the dense phase and the dilute phase are considered separately. This required distinguishing the two phases. The dense phase and dilute phase were determined by density analysis. To this end, a density profile was measured every 10 ns from snapshots such as those shown in Fig. 2a and b. For each time point, the density profile was shifted to put the center of mass of Rubisco molecules at the center of the box, which centered the dense phase. Subsequently, the “dense phase” was defined as the central region away from interface with the dilute phase – in practice, we chose the region where the average Rubisco density was at least 80% of its maximum. Similarly, the “dilute phase” was defined as the flanking regions away from the interface. Because of the larger region of dilute phase, we included only zones well away from the interface, where the average density is flat within numerical noise.

To enumerate the complexes found in the dilute phase as shown in Fig. 3c, we employed a custom cluster code^49^. The cluster code computes molecular bonds between Rubiscos and EPYC1s via the bonding criteria described in the section Dissociation constant 𝐾_d_.

From the same definition of a bond, the number of satisfied stickers was enumerated for each Rubisco in both the dense and dilute phases, along with the number of EPYC1 molecular partners per Rubisco, as shown in Fig. 3b and d.

### Dimer-gel theory

We consider a dimer-gel theory which has three terms (see Refs. 11 and 35):

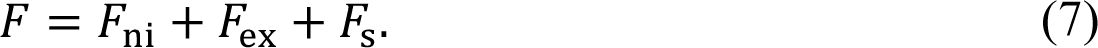

The first term considers non-interacting, ideal, contributions to the free energy given by

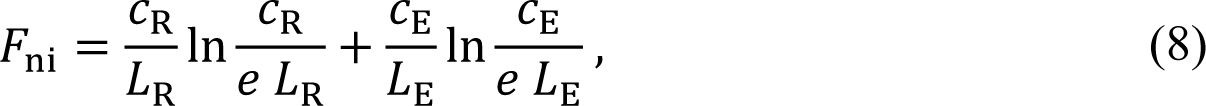

where 𝑐_R_ and 𝑐_E_ are the sticker concentrations for Rubisco and EPYC1, and 𝐿_*R*_ = 8 and 𝐿_E_ = 5 are the number of stickers for a single Rubisco and single EPYC1, respectively. The second term 𝐹_ex_ accounts for excluded volume interactions among the molecules, which we take to be hard spheres:

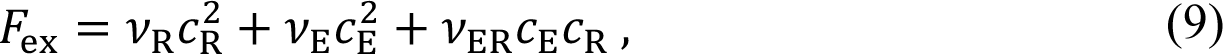

where the effective volume constants 𝜈_R_, 𝜈_E_, and 𝜈_ER_ are obtained by a virial expansion^50, 51^. The Rubisco-Rubisco effective volume is four times an eighth of its molecular volume with diameter 𝑑 = 10 nm given by 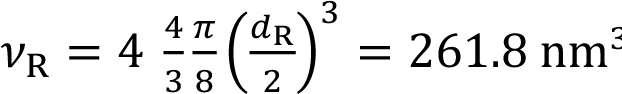. The EPYC1-EPYC1 effective volume is 4 times its effective sticker volume, where a sticker for EPYC1 consists of approximately 60 amino acids that are each 3-4 Å in length, which includes the length of the region responsible for binding Rubisco as well as a linker^52^, in terms of its radius of gyration *R_g_* ≈ 1 given by 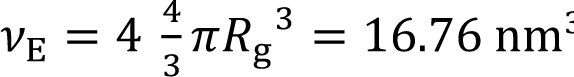. The effective Rubisco-EPYC1 volume is determined by the effective radius of the two such that 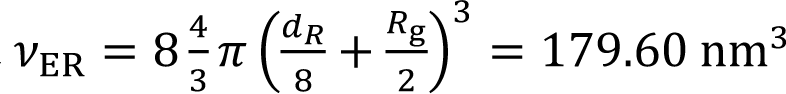. These volumes are written in terms of molarity which involves multiplication by Avogadro’s number after being converted to liters.

The final term, 𝐹_*s*_, which describes sticker-sticker bonding is taken to be the minimum of the free energy to form molecular dimers or the free energy to form independent sticker-sticker pairs (since we find one of these free energies always dominates the other),

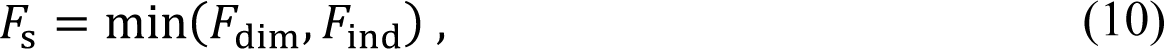

with

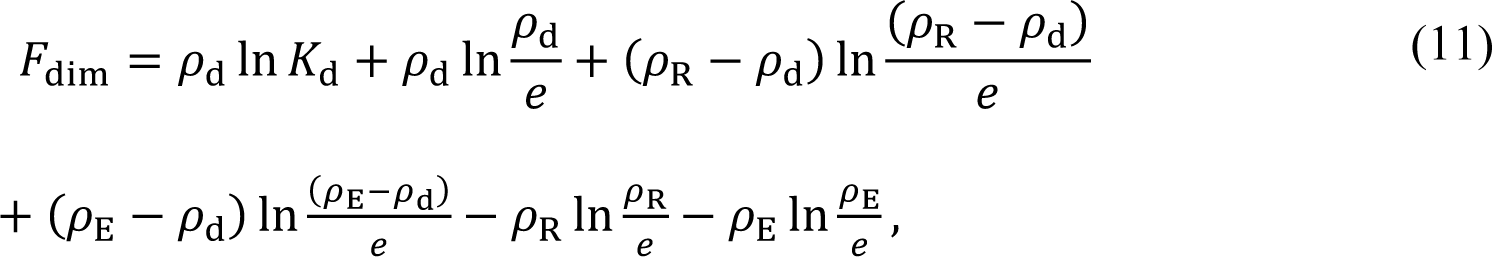

and

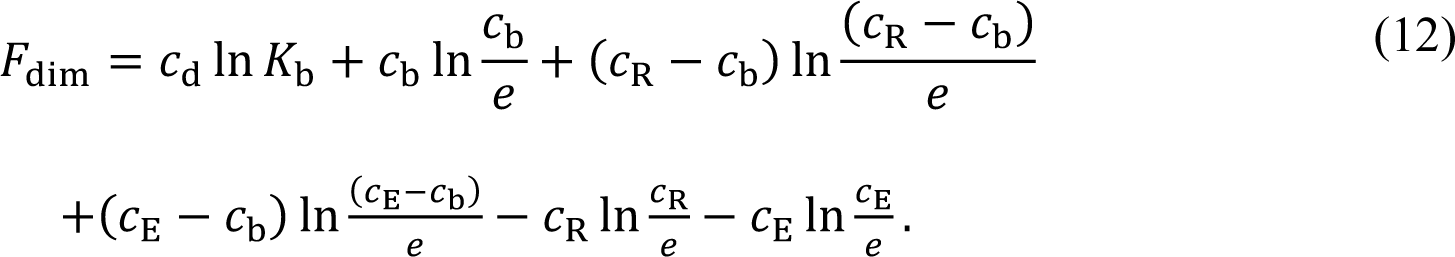

Here, 𝜌_R_ and 𝜌_E_ are the molecular concentrations of Rubisco and EPYC1, 𝑐_R_ and 𝑐_E_ are the respective sticker concentrations of Rubisco and EPYC1, and 𝜌_d_ and 𝑐_b_ are the concentrations of molecular dimers and independent sticker-sticker pairs given, respectively, by the following:

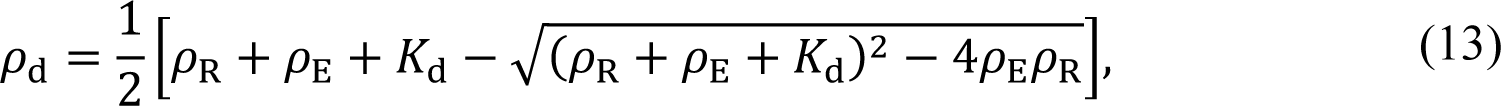

and

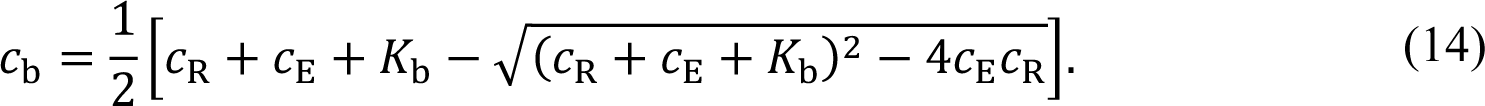

The convex hull of the total free energy 𝐹 determines the phase diagram. In practice, the only additional inputs required by the dimer-gel theory are the molecular dissociation constant, 𝐾_d_, and the sticker-sticker dissociation constant, 𝐾_b_. In Fig. 4, we use the dissociation constant 𝐾_d_ from Fig. 3a that corresponds to 𝑈_0_ = 14 𝑘_B_𝑇 for each of the two linker lengths. The 𝐾_b_ value is fit separately for each linker length so that the dilute-phase concentrations within the dimer-gel theory match the simulation results shown in Fig. 2.

### Rod-like order parameter

The transition to a gas of rods is quantified by the alignment of Rubisco molecules given by the average cosine of the relative angle between all pairs of Rubisco that are neighbors of the 𝑖’th Rubisco,

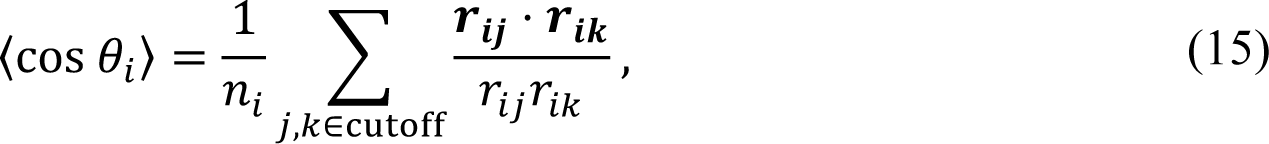

where bold 𝒓_𝒊𝒋_ and 𝒓_𝒊𝒌_ are, respectively, the vectors from the center of Rubisco 𝑖 to the centers of Rubiscos 𝑗 and 𝑘, for all neighbors whose absolute distances 𝑟_*ij*_ and 𝑟_*ik*_ fall within a cutoff distance of 15 nm to include only nearest neighbors. 𝑛_*i*_ is the number Rubisco pairs enumerated in the sum. This observable is then averaged over all Rubisco molecules to get

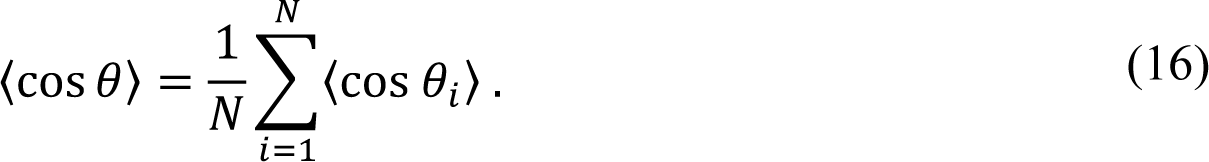

### Conditional probability to bond two consecutive stickers in a two-sticker system

To understand why bonded configurations of EPYC1 and Rubisco depend on EPYC1 linker length, we first consider a simpler system of a truncated EPYC1 molecule (EPYC12) consisting of two stickers connected by a linker. There is no excluded volume between the stickers and the linker is modeled as a harmonic spring with energy

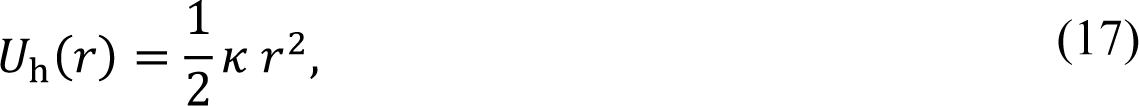

where 𝑟 is the relative distance between the two stickers, and 𝜅 is the spring constant. For this system, the equilibrium effective linker length is given by

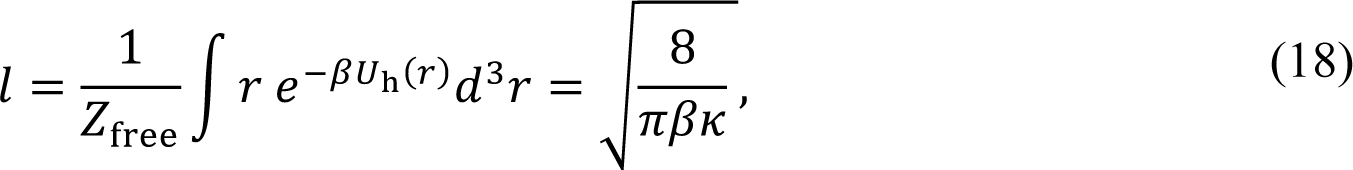

where 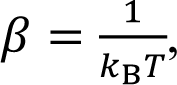, and

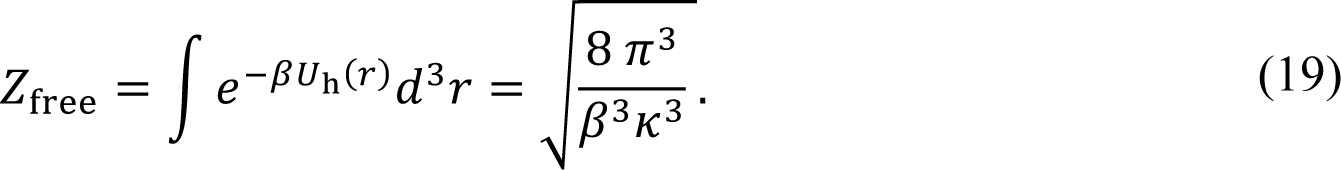

From Eq. (18), we can relate the spring constant to the effective linker length,

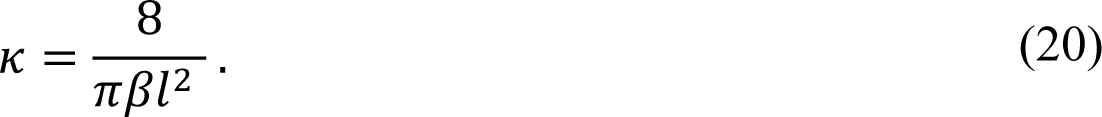

Now consider two stickers for the EPYC12 stickers that are a distance 𝑑 apart, with one of the stickers at 𝒓_𝟏_ = (0,0,0) and the other at 𝒓_𝟐_ = (𝑑, 0,0). Given that one sticker of EPYC12 is bound at exactly 𝒓_𝟏_,we want to calculate the conditional probability that the other sticker is bound at 𝒓_𝟐_. At thermal equilibrium, this conditional probability is given by

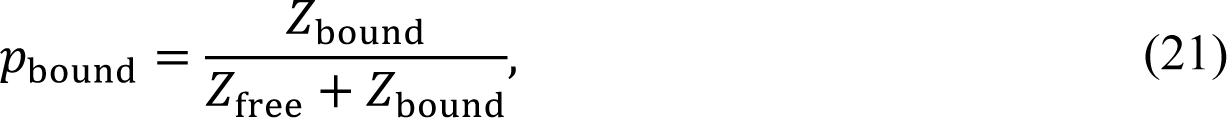

where 𝑍_bound_ is the partition function of the second sticker being bound at a distance, 𝑑,

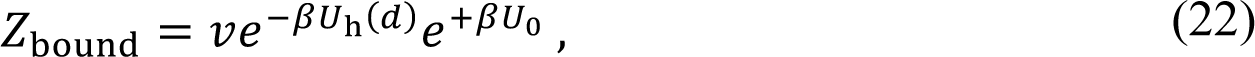

where 𝑣 is the volume over which a bond can be made, and 𝑈_0_ is the bonding energy. More generally, we can express 𝑍_bound_ in terms of the sticker dissociation constant, 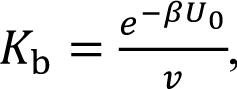, so that

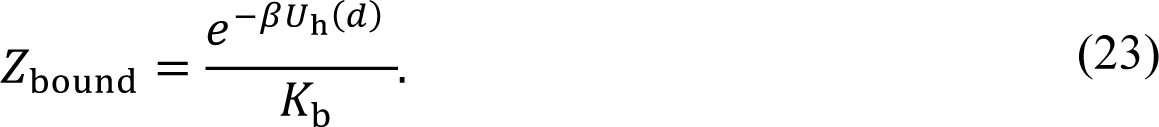

Thus, the conditional probability for the second sticker to be bound is

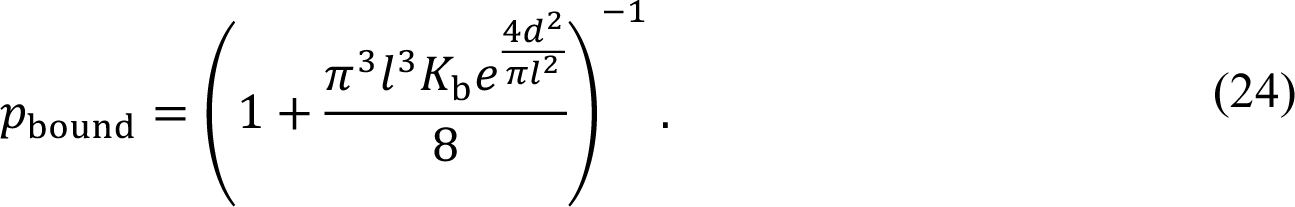

The result in Eq. (24) implies that it is unfavorable for two adjacent EPYC1 stickers to bind neighboring Rubisco stickers if the effective EPYC1 linker length 𝑙 is substantially smaller than the sticker spacing 𝑑. This has strong implications for the observed of Rubisco-EPYC1 bonding arrangements in both dense and dilute phases in our simulations.

### Estimation of linker lengths and Rubisco sticker spacing in carboxysomes

Approximating a disordered protein as an ideal chain, the radius of gyration is given by 𝑅_g_ = 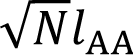, where 𝑁 is the number of amino acids in the protein and 𝑙_AA_ ≈ 0.39 nm is the length of an amino acid^52^. For a protein composed of stickers and linkers, the effective linker length is then given by 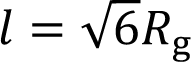 in terms of a single linker’s radius of gyration (see Supplementary Note 1). If we approximate CsoS2 linkers as 50 AAs, this gives a radius of gyration of 𝑅_g_ = 2.8 nm and an effective linker length of 𝑙 = 6.8 nm. Likewise, CcmM linkers are about 30 AAs, which gives a radius of gyration of 𝑅_g_ = 2.1 nm and an effective linker length of 𝑙 = 5.2 nm.

Next, we would like to estimate the distance between Rubisco stickers for alpha and beta cyanobacteria. Given that Rubisco has a diameter of 12 nm, its circumference is 𝐶 = 2𝜋 (6 nm)∼ 37.7 nm. For alpha cyanobacteria, the spacing between adjacent sets of north-south sticker pairs is therefore 𝐶⁄4 = 9.4 nm, and the distance between the individual Rubisco stickers in beta cyanobacteria is the same.

## Acknowledgements

We thank Ariel Amir, Guanhua He, Shan He, Wencheng Ji, Ofer Kimchi, Linnea Lemma, Sam Safran, David Sivak, Jon Yuly for useful discussions. This work was supported by grants from the National Institutes of Health (R01GM140032), the National Science Foundation (MCB-1935444), and the National Science Foundation through the Center for the Physics of Biological Function (PHY-1734030). T.G. was supported by the Schmidt Science Fellowship.

## Author contributions

T.G., Y.Z, M.C.J and N.S.W. designed the study. T.G. performed all simulations and analytical calculations. T.G. and N.S.W. developed theory. All authors contributed to interpreting results and preparing the manuscript.

## Data availability

Data or codes supporting the findings of this manuscript are available from the corresponding author upon reasonable request.

