## Supplemental Information for "Effects of linker length on phase separation: lessons from the Rubisco-EPYC1 system of the algal pyrenoid"

### Supplementary Note 1

#### All-atom simulations:

The long EPYC1 linkers of the coarse-grained model were parametrized via comparison with an all-atom simulation of EPYC1. The EPYC1 molecule contains 4531 atoms and a net positive charge of +29. All-atom simulations of EPYC1 were performed using Desmond software[1,2] with explicit solvent with 180,657 water molecules, 327 sodium ions Na+, and 356 chloride ions Cl- in a cubic box of 17.6 nm edge length. The simulations were first equilibrated, and then measurements were taken over a total time of 5 μs. In Supplementary Fig. 1, the radius of gyration is shown in increments of 0.1 μs.

From these all-atom trajectories, the average radius of gyration is $\langle R_{\mathrm{g}} \rangle = 3.9 \pm 0.7$ nm. By approximating EPYC1 as a ideal linear chain, the total average radius of gyration can be written in terms of the equilibrium end-to-end length of a linker, $l$, and the number of linkers in the chain[3], $n$, as $\langle R_{\mathrm{g}}^2 \rangle = nl^2/6$. Using the all-atom average radius of gyration $\langle R_{\mathrm{g}} \rangle =$ 3.9 nm and $n = 4$, the average end-to-end distance for an EPYC1 linker is $l = 4.8 \pm 0.9$ nm. We use this value of $l$ to fit the average sticker-sticker distance of a single EPYC1 molecule in the coarse-grained simulations, yielding a spring constant $k = 0.12\ k_{\mathrm{B}}T/\sigma^2$, where $\sigma = 2$ nm is the sticker diameter. Note that this is these linkers are referred to as the long "linkers" in the main text.

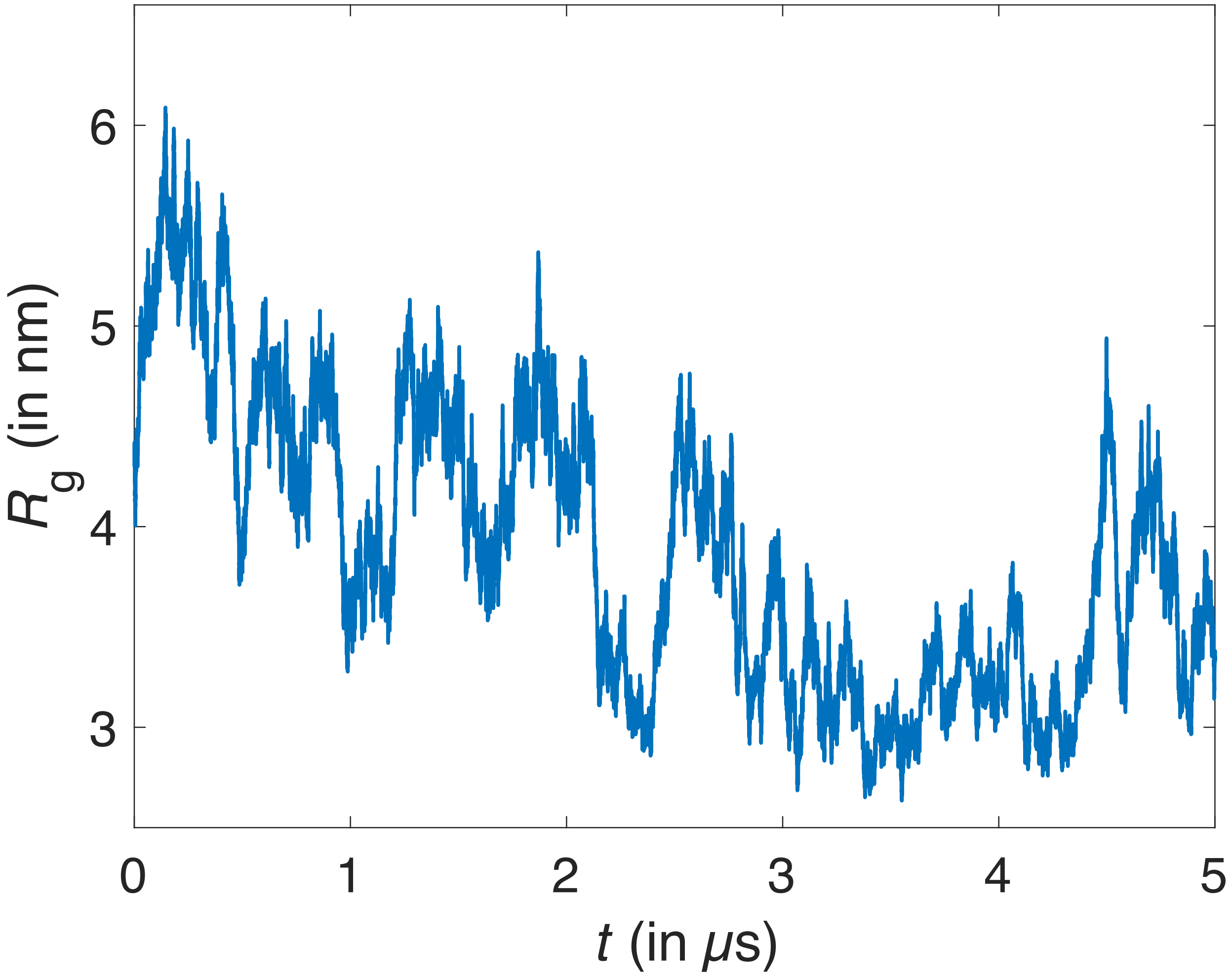


Supplementary Figure 1. EPYC1 radius of gyration as a function of time from all-atom simulations. The force fields for the all-atom simulation are the same as in Ref. 2.

**Supplementary Note 2**

**High EPYC1 to Rubisco relative stoichiometries:**

Supplementary Fig. 2a and b show representative simulation snapshots for 3:1 and 10:1 EPYC1:Rubisco sticker stoichiometry for long EPYC1 linkers. Corresponding snapshots for short linkers are shown in Supplementary Fig. 2c and d. The short EPYC1 linkers still mediate phase separation even at these high relative stoichiometries, but the long linkers do not. These results indicate that the 2:1 EPYC1:Rubisco stoichiometry in Figs. 2 and 3 in the main text is near the phase boundary for long linkers, but well within the two-phase region for short linkers.

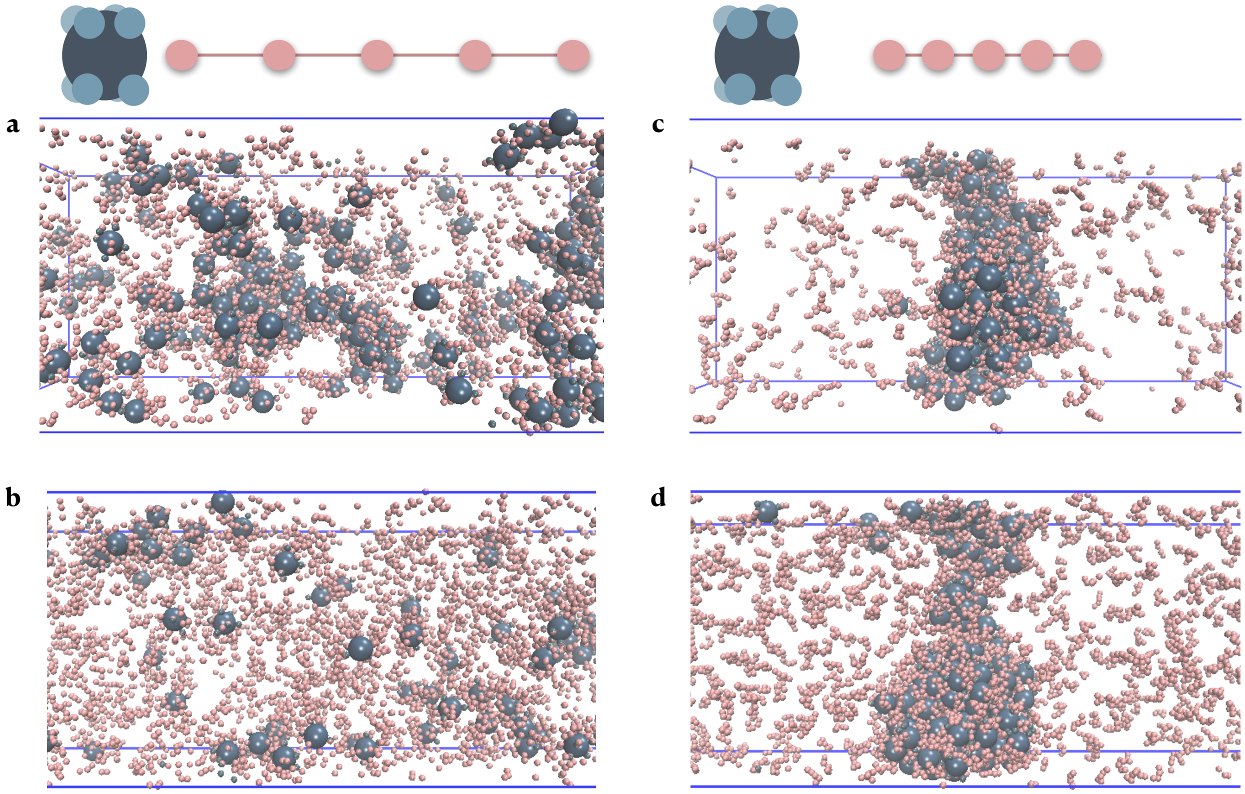


Supplementary Figure 2. Snapshots from simulations of Rubisco-EPYC1 systems with the same parameters as in Fig. 2 at different stoichiometries and overall polymer concentrations. **a,b** Long EPYC1 linkers at EPYC1:Rubisco sticker stoichiometry of 3:1 (864 EPYC1s, 180 Rubiscos in a box of size 315 nm x 126 nm x 126 nm with periodic boundaries) **a,** and 10:1 (2080 EPYC1s, 130 Rubiscos in a box of size 630 nm x 126 nm x 126 nm with periodic boundaries) **b**. **c,d** as in **a,b** but for short EPYC1 linkers.

**Transition to gas of rods at 1:1 EPYC1 to Rubisco sticker stoichiometry:**

Here, we show that the transition from a phase-separated state to a gas of rods as a function of Rubisco sticker position is robust with respect to sticker stoichiometry. In the main text, Fig. 5d shows that the gas of rods forms for short EPYC1 linkers at 2:1 EPYC1 to Rubisco sticker stoichiometry. Supplementary Fig. 3 shows that there is also a transition to a gas of rods at 1:1 sticker stoichiometry.

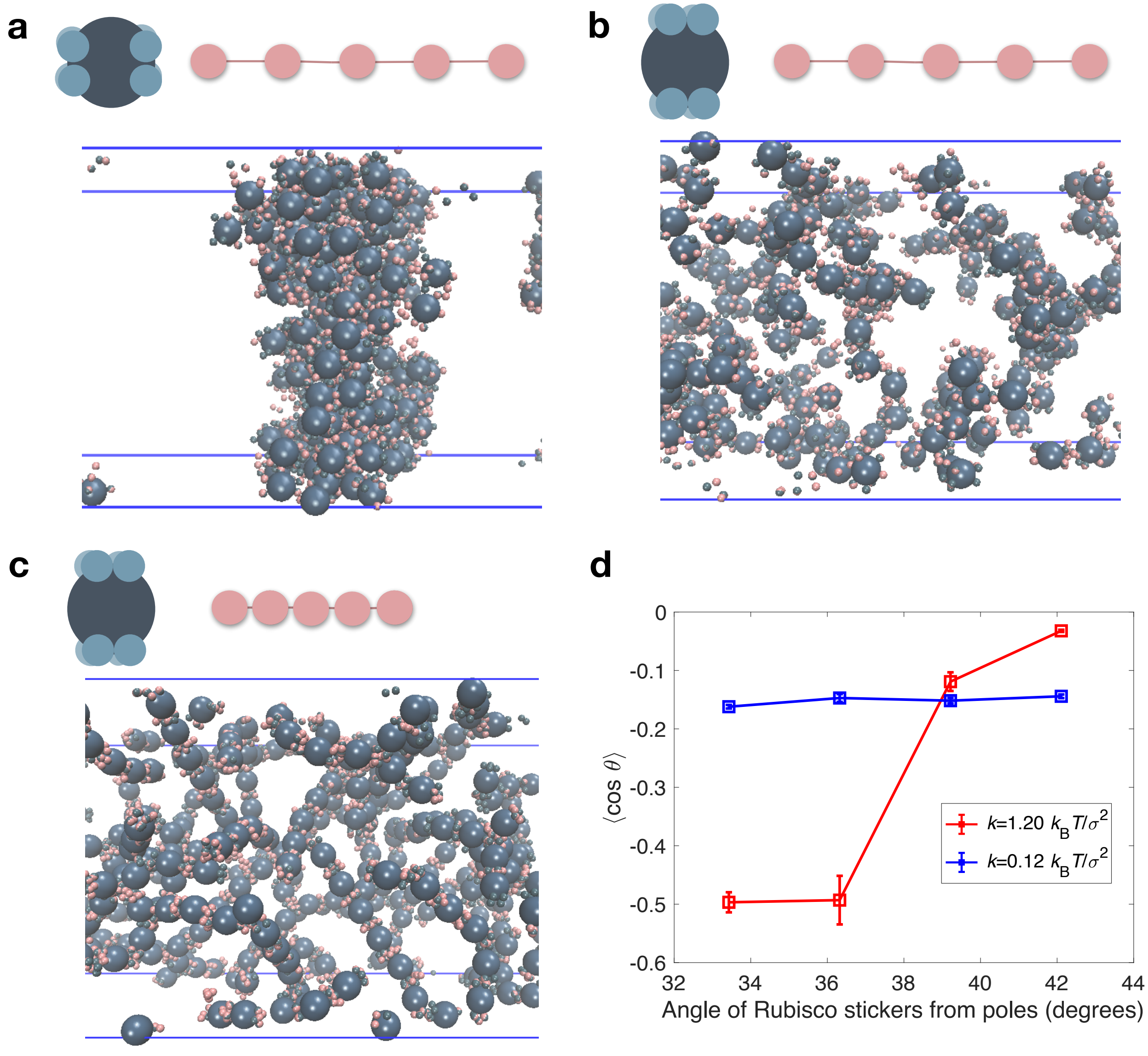


Supplementary Figure 3. A transition to rods as a function of Rubisco sticker location. Rubisco-EPYC1 systems with long ($k = 0.12 \frac{k_B T}{\sigma^2}$) or short EPYC1 linkers ($k = 1.20 \frac{k_B T}{\sigma^2}$) were simulated at equal sticker stoichiometry (608 EPYC1s and 380 Rubiscos) for $U_0 = 14\, k_B T$ in boxes of size 315 nm x 126 nm x 126 nm with periodic boundaries at a temperature $T = 300$ K. **a** Snapshot for Rubisco stickers located at 79 degrees from the poles with long EPYC1 linkers shows phase separation. **b** Snapshot for Rubisco stickers located at 33 degrees from the poles with long

EPYC1 linkers shows that instead of phase separating, the system forms a gas of small complexes. **c** Snapshot for Rubisco stickers located at 33 degrees from the poles with short EPYC1 linkers shows that the system forms a gas rods. **d** Order parameter $\langle \cos \theta \rangle$ for rods as a function of angle of Rubisco stickers from poles, where $\theta$ is the angle formed by neighboring Rubiscos around a central Rubisco (see Methods for details). The blue symbols are for long linkers and the red symbols are for short linkers. Error bars are SD values obtained from three independent simulations.

**Rubisco with movable stickers:**

To better understand the inverse relationship between robust phase separation and the molecular fit between EPYC1 linker length and Rubisco sticker spacing, we allowed Rubisco stickers to freely diffuse on the surface of the 12 nm diameter base instead of keeping them at fixed locations. All of the equations are the same as those noted in the section Model details, but now the Rubisco stickers are held to the base with a harmonic bond given by

$$U_{\mathrm{R}}(r) = \frac{1}{2} k_{\mathrm{R}} (r - r_0)^2, \tag{S1}$$

where $r_0 = 6$ nm and $k_{\mathrm{R}} = 19\, k_{\mathrm{B}} T/\sigma^2$. The spring is held tight by the large value of $k_{\mathrm{R}}$ to allow for little motion away from the surface of the base.

The molecular fit is quantified by the Rubisco-EPYC1 molecular dissociation constant. At a value of $U_0 = 14\, k_{\mathrm{B}} T$, we find that the short-linker dissociation constant is $K_{\mathrm{d}} = 74 \pm 20$ µM which is about four times smaller than the short-linker dissociation constant shown in Fig. 3a, i.e. the free movement of the Rubisco stickers improves the molecular fit. We would expect that if diffusing stickers improves molecular fit, that this system would not phase separate well. In Supplementary Fig. 4, we show a snapshot of the system with movable Rubisco stickers and with the same parameters as in the main text for the short EPYC1 linkers. As expected, the molecular fit is greatly improved with one EPYC1 preferring to be bound to one Rubisco instead of spanning multiple Rubiscos.

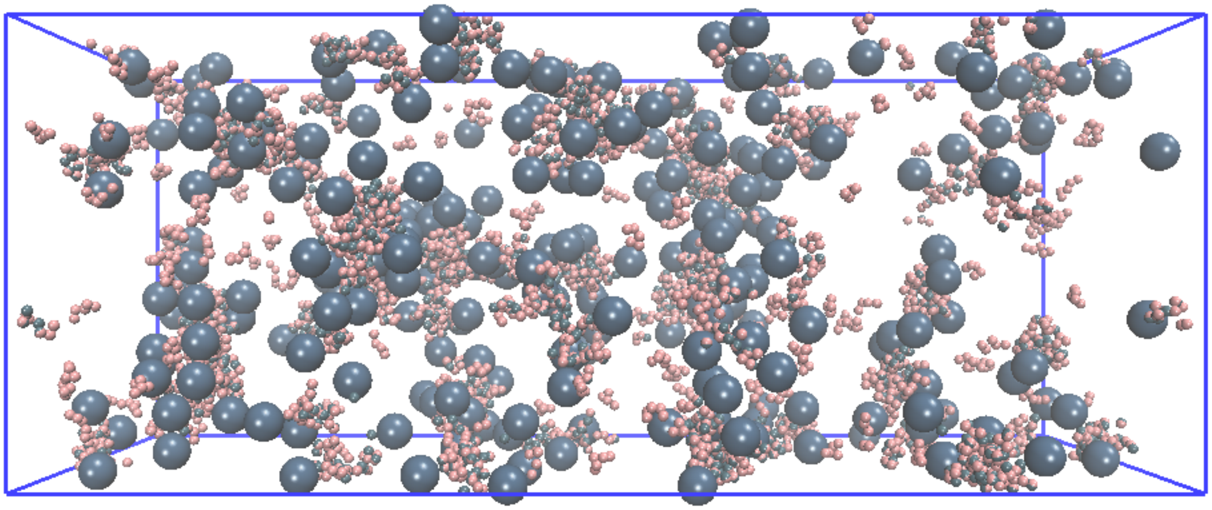

Supplementary Figure 4. A Rubisco-EPYC1 system with Rubisco stickers that are free to move on the surface of the spherical Rubisco base. A snapshot is shown for short EPYC1 linkers at 2:1 EPYC1 to Rubisco sticker stoichiometry (760 EPYC1s, 240 Rubiscos in a box of size 315 nm x 126 nm x 126 nm with periodic boundaries).
